# Normalization of the microbiota in patients after treatment for colonic lesions

**DOI:** 10.1101/138222

**Authors:** Marc A Sze, Nielson T Baxter, Mack T Ruffin, Mary AM Rogers, Patrick D Schloss

## Abstract

**Background:** Colorectal cancer is a worldwide health problem. Despite growing evidence that members of the gut microbiota can drive tumorigenesis, little is known about what happens to it after treatment for an adenoma or carcinoma. This study tested the hypothesis that treatment for adenoma or carcinoma alters the abundance of bacterial populations associated with disease to those associated with a normal colon. We tested this hypothesis by sequencing the 16S rRNA genes in the feces of 67 individuals before and after treatment for adenoma (N = 22), advanced adenoma (N = 19), and carcinoma (N = 26).

**Results:** There were small changes to the bacterial community associated with adenoma or advanced adenoma and large changes associated with carcinoma. The communities from patients with carcinomas changed significantly more than those with adenoma following treatment (P-value < 0.001). Although treatment was associated with intrapersonal changes, the change in the abundance of individual OTUs in response to treatment was not consistent within diagnosis groups (P-value > 0.05). Because the distribution of OTUs across patients and diagnosis groups was irregular, we used the Random Forest machine learning algorithm to identify groups of OTUs that could be used to classify pre and post-treatment samples for each of the diagnosis groups. Although the adenoma and carcinoma models could reliably differentiate between the pre and post-treatment samples (P-value < 0.001), the advanced-adenoma model could not (P-value = 0.61). Furthermore, there was little overlap between the OTUs that were indicative of each treatment. To determine whether individuals who underwent treatment were more likely to have OTUs associated with normal colons we used a larger cohort that contained individuals with normal colons and those with adenomas, advanced adenomas, and carcinomas. We again built Random Forest models and measured the change in the positive probability of having one of the three diagnoses to assess whether the post-treatment samples received the same classification as the pre-treatment samples.

Samples from patients who had carcinomas changed towards a microbial milieu that resembles the normal colon after treatment (P-value < 0.001). Finally, we were unable to detect any significant differences in the microbiota of individuals treated with surgery alone and those treated with chemotherapy or chemotherapy and radiation (P-value > 0.05).

**Conclusions:** By better understanding the response of the microbiota to treatment for adenomas and carcinomas, it is likely that biomarkers will eventually be validated that can be used to quantify the risk of recurrence and the likelihood of survival. Although it was difficult to identify significant differences between pre and post-treatment samples from patients with adenoma and advanced adenoma, this was not the case for carcinomas. Not only were there large changes in pre versus post-treatment samples for those with carcinoma, but these changes were towards a more normal microbiota.

## Background

Colorectal cancer (CRC) is the third most common cause of cancer deaths in the United States (1, 2). Disease mortality has significantly decreased, predominately due to improvements in screening (2). Despite these improvements, there are still approximately 50,000 CRC-related deaths per year in the United States (1). Current estimates indicate that 20-30% of those who undergo treatment will experience recurrence and 35% of all patients will die within five years (3–5). Identification of methods to assess patients’ risk of recurrence is of great importance to reduce mortality and healthcare costs.

There is growing evidence that the gut microbiota is involved in the progression of CRC. Mouse-based studies have identified populations of *Bacteroides fragilis, Escherichia coli*, and *Fusobacterium nucleatum* that alter disease progression (6–10). Furthermore, studies that shift the structure of the microbiota through the use of antibiotics or inoculation of germ free mice with human feces have shown that varying community compositions can result in varied tumor burden (11–13). Collectively, these studies support the hypothesis that the microbiota can alter the amount of inflammation in the colon and with it the rate of tumorigenesis (14).

Building upon this evidence, several human studies have identified unique signatures of colonic lesions (15–20). One line of research has identified community-level differences between those bacteria that are found on and adjacent to colonic lesions and have supported a role for *Bacteroides fragilis, Escherichia coli*, and *Fusobacterium nucleatum* in tumorigenesis (21–23). Others have proposed feces-based biomarkers that could be used to diagnose the presence of colonic adenomas and carcinomas (24–26). These studies have associated Fusobacterium nucleatum and other oral pathogens with colonic lesions (adenoma, advanced adenoma, and carcinoma). They have also noted that the loss of bacteria generally thought to produce short chain fatty acids, which can suppress inflammation, is associated with colonic lesions. This suggests that gut bacteria have a role in tumorigenesis with potential as useful biomarkers for aiding in the early detection of disease (21–26).

Despite advances in understanding the role between the gut microbiota and colonic tumorigenesis, we still do not understand how treatments including resection, chemotherapy, and/or radiation affect the composition of the gut microbiota. If the microbial community drives tumorigenesis then one would hypothesize that treatment to remove a lesion would not only remove the lesion, but also the microbiota that promoted the tumorigenesis and hence the risk of recurrence. To test this hypothesis, we addressed two related questions: Does treatment affect the colonic microbiota in a predictable manner? If so, does the treatment alter the community to more closely resemble that of individuals with normal colons?

We answered these questions by sequencing the V4 region of 16S rRNA genes amplified from fecal samples of individuals with adenoma, advanced adenoma, and carcinomas pre and post-treatment. We used classical community analysis to compare the alpha and beta-diversity of communities pre and post-treatment. Next, we generated Random Forest models to identify bacterial populations that were indicative of treatment for each diagnosis group. Finally, we measured the predictive probabilities to assess whether treatment yielded bacterial communities similar to those individuals with normal colons. We found that treatment alters the composition of the gut microbiota and that, for those with carcinomas, the gut microbiota shifted more towards that of a normal colon after treatment. In the individuals with carcinomas, no difference was found by the type of treatment (surgery alone, surgery with chemotherapy, surgery with chemotherapy and radiation). Understanding how the community responds to these treatments could be a valuable tool for identifying biomarkers to quantify the risk of recurrence and the likelihood of survival.

## Results

### Treatment for colonic lesions alters the bacterial community structure

Within our 67-person cohort we tested whether the microbiota of patients with adenoma (N = 22), advanced adenoma (N = 19), or carcinoma (N = 26) had any broad differences between pre and post-treatment samples [Table 1]. None of the individuals in this study had any recorded antibiotic usage that was not associated with surgical treatment of their respective lesion. The structure of the microbial communities of the pre and post-treatment samples differed, as measured by the *θ*YC beta diversity metric [Figure 1A]. We found that the communities obtained pre and post-treatment among the patients with carcinomas changed significantly more than those patients with adenoma (P-value < 0.001). There were no significant differences in the amount of change observed between the patients with adenoma and advanced adenoma or between the patients with advanced adenoma and carcinoma (P-value > 0.05). Next, we tested whether there was a consistent direction in the change in the community structure between the pre and post-treatment samples for each of the diagnosis groups [Figure 1B-D]. We only observed a consistent shift in community structure for the patients with carcinoma when using a PERMANOVA test (adenoma P-value = 0.999, advanced adenoma P-value = 0.945, and carcinoma P-value = 0.005). Finally, we measured the number of observed OTUs, Shannon evenness, and Shannon diversity in the pre and post-treatment samples and did not observe a significant change for any of the diagnosis groups (P-value > 0.05) [Table S1].

**Table 1:** Demographic data of patients in the pre and post-treatment cohort

|  | Adenoma | Advanced Adenoma | Carcinoma |
| --- | --- | --- | --- |
| n | 22 | 19 | 26 |
| Age (Mean ± SD) | 61.68 ± 7.2 | 63.11 ± 10.9 | 61.65 ± 12.9 |
| Sex (%F) | 36.36 | 36.84 | 42.31 |
| BMI (Mean ± SD) | 26.86 ± 3.9 | 25.81 ± 4.7 | 28.63 ± 7.2 |
| Caucasian (%) | 95.45 | 84.21 | 96.15 |
| Days Between Colonoscopy (Mean ± SD) | 255.41 ± 42 | 250.16 ± 41 | 350.85 ± 102 |
| Surgery Only | 4 | 4 | 12 |
| Surgery & Chemotherapy | 0 | 0 | 9 |
| Surgery, Chemotherapy, & Radiation | 0 | 0 | 5 |

**Figure 1:**
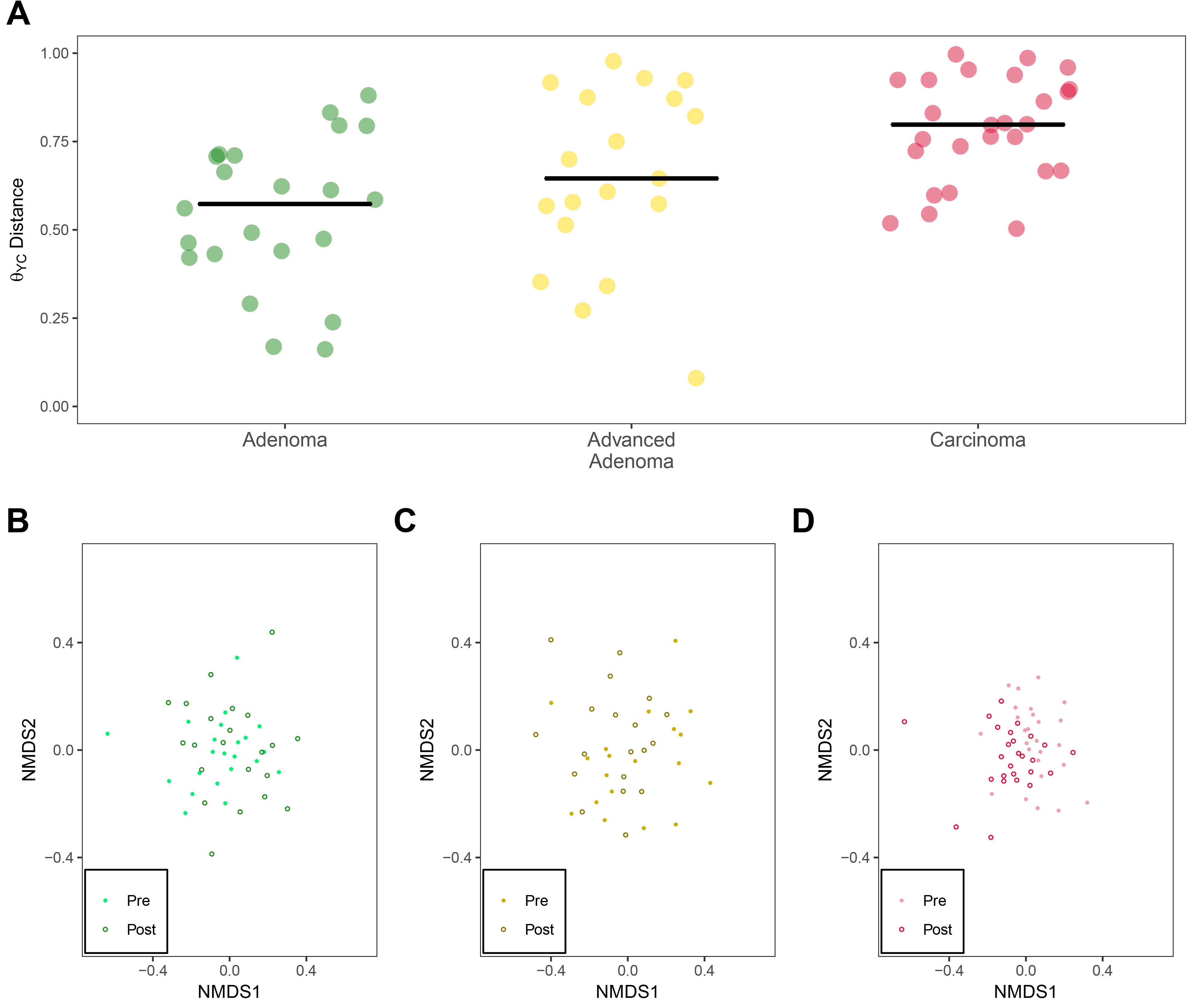
General differences between adenoma, advanced adenoma, and carcinoma groups after treatment. **A)** *θ*YC distances from pre versus post sample within each individual. A significant difference was found between the adenoma and carcinoma group (P-value = 5.36e-05). Solid black points represent the median value for each diagnosis group. B) NMDS of the pre and post-treatment samples for the adenoma group. C) NMDS of the pre and post-treatment samples for the advanced adenoma group. D) NMDS of the pre and post-treatment samples for the carcinoma group.

### The treatment of lesions are not consistent across diagnosis groups

We used two approaches to identify those bacterial populations that change between the two samples for each diagnosis group. First, we sought to identify individual OTUs that could account for the change in overall community structure. However, using a paired Wilcoxon test we were unable to identify any OTUs that were significantly different in the pre and post-treatment groups (P-value > 0.05). It is likely that high inter-individual variation and the irregular distribution of OTUs across individuals limited the statistical power of the test. We attempted to overcome these problems by using Random Forest models to identify collections of OTUs that would allow us to differentiate between pre and post-treatment samples from each of the diagnosis groups. The adenoma and carcinoma models performed well (adenoma AUC range = 0.54 - 0.83 and carcinoma AUC range = 0.82 - 0.98); however, the model for patients treated for advanced adenomas was not able to reliably differentiate between the pre and post-treatment samples (advanced adenoma AUC range = 0.34 - 0.65). Interestingly, the top 10 most important OTUs by MDA that were used for each model had little overlap with each other [Figure 2]. Although treatment had an impact on the overall community structure, the effect of treatment was not consistent across patients and diagnosis groups. Both the adenoma and carcioma treatment models had AUCs that were significantly higher than a random model permutation (P-value < 0.0001).

**Figure 2:**
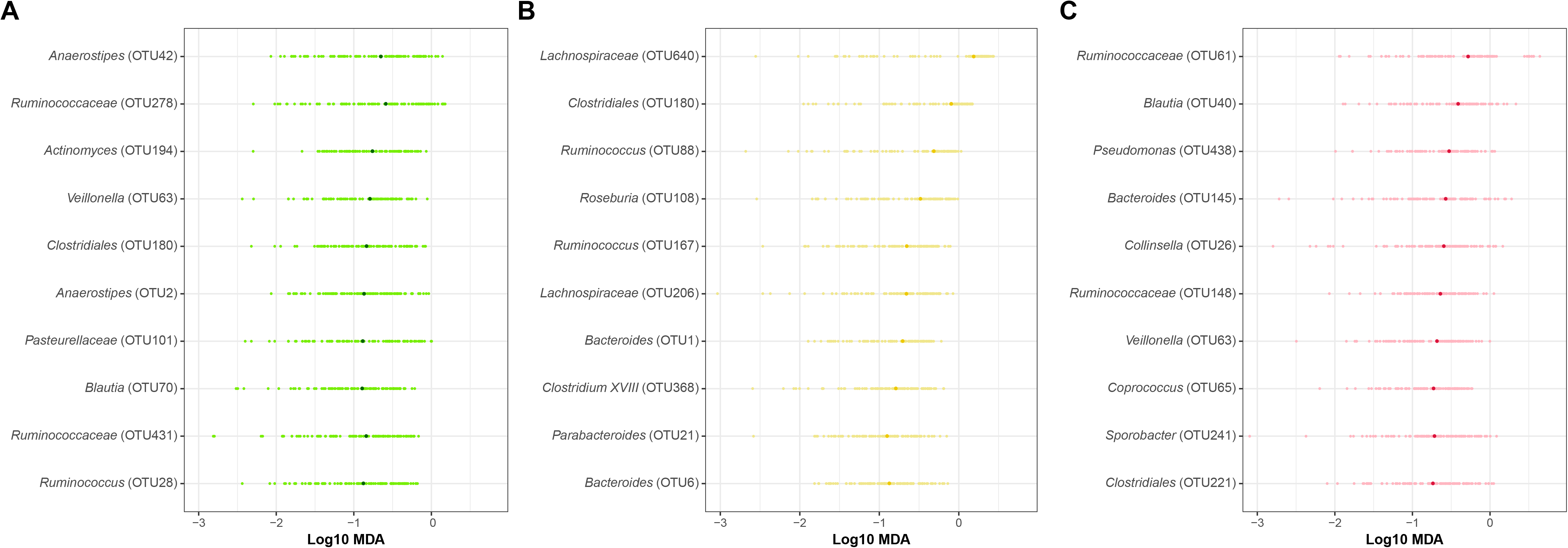
The top 10 most important OTUs used to classify treatment for adenoma, advanced adenoma, and carcinoma. **A)** Adenoma OTUs. B) Advanced Adenoma OTUs. C) Carcinoma OTUs. The darker circle highlights the median log10 MDA value obtained from 100 different 80/20 splits while the lighter colored circles represents the value obtained for a specific run.

### Post-treatment samples from patients with carcinoma more closely resemble those of a normal colon

Next, we determined whether treatment changed the microbiota in a way that the post-treatment communities resembled that of patients with normal colons. To test this, we used an expanded cohort of 423 individuals that were diagnosed under the same protocol as having normal colons or colons with adenoma, advanced adenoma, or carcinoma [Table 2]. We then constructed Random Forest models to classify the study samples, with the 3 diagnosis groups (adenoma, advanced adenoma, or carcinoma), or having a normal colon. The models performed moderately with CRC being the best (adenoma AUC range = 0.50 - 0.62, advanced adenoma AUC range = 0.53 - 0.67, carcinoma AUC range = 0.71 - 0.82; Figure S1). The OTUs that were in the top 10% of importance for the adenoma and advanced adenoma models largely overlapped and those OTUs that were used to classify the carcinoma samples were largely distinct from those of the other two models [Figure 3A]. Among the OTUs that were shared across the three models were those populations generally considered beneficial to their host (e.g. Faecalibacterium, Lachnospiraceae, Bacteroides, Dorea, Anaerostipes, and Roseburia) [Figures 3B]. Although many of important OTUs in the top 10% were also included in the model differentiating between patients with normal colons and those with carcinoma, this model also included OTUs affiliated with populations that have previously been associated with carcinoma (Fusobacterium, Porphyromonas, Parvimonas) (24–26) [Figure S2] with some individuals showing a marked decrease in relative abundance [Figure S3]. Finally, we applied these three models to the pre and post-treatment samples for each diagnosis group and quantified the change in the positive probability of the model. A decrease in the positive probability would indicate that the microbiota more closely resembled that of a patient with a normal colon. There was no significant change in the positive probability for the adenoma or advanced adenoma groups (P-value > 0.05) [Figure 4]. The positive probability for the pre and post-treatment samples from patients diagnosed with carcinoma significantly decreased with treatment, suggesting a shift toward a normal microbiota for most individuals (P-value = 0.001). Only, 7 of the 26 patients (26.92%) who were diagnosed with a carcinoma had a higher positive probability after treatment; one of those was re-diagnosed with carcinoma on the follow up visit. These results indicate that, although there were changes in the microbiota associated with treatment, those experienced by patients with carcinoma after treatment yielded gut bacterial communities of greater similarity to that of a normal colon.

**Table 2:** Demographic data of training cohort

|  | Normal | Adenoma | Advanced Adenoma | Carcinoma |
| --- | --- | --- | --- | --- |
| n | 172 | 67 | 90 | 94 |
| Age (Mean ± SD) | 54.29 ± 9.9 | 63.01 ± 13.1 | 64.07 ± 11.3 | 64.37 ± 12.9 |
| Sex (%F) | 64.53 | 46.27 | 37.78 | 43.62 |
| BMI (Mean ± SD) | 26.97 ± 5.3 | 25.69 ± 4.8 | 26.66 ± 4.9 | 29.27 ± 6.7 |
| Caucasian (%) | 87.79 | 92.54 | 92.22 | 94.68 |

**Figure 3:**
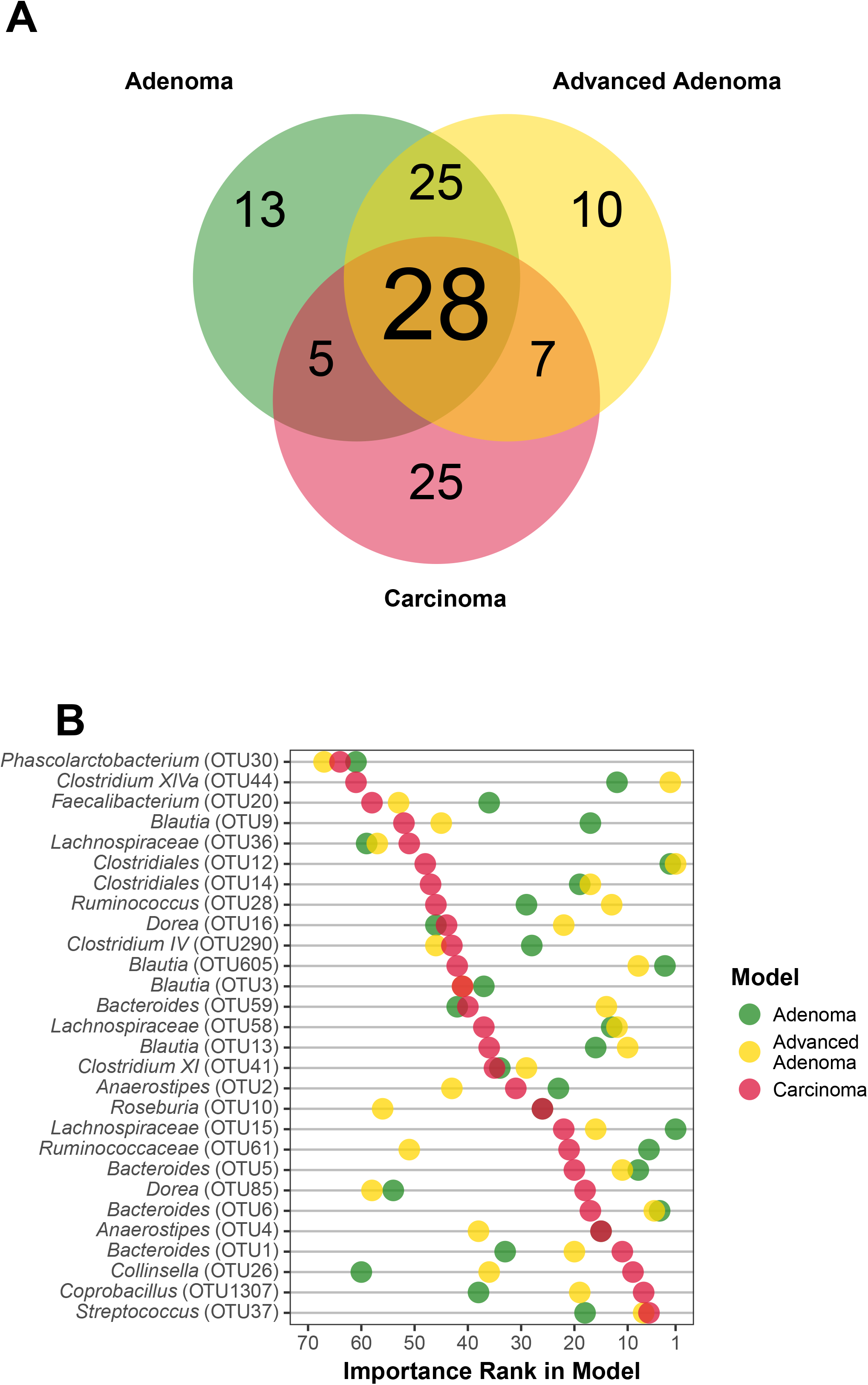
Top 10% most important OTUs common to those models used to differentiate between patients with normal colons and those with adenoma, advanced adenoma, and carcinoma. **A)** Venn diagram showing the OTU overlap between each model. B) For each common OTU the lowest taxonomic identification and importance rank for each model run is shown.

**Figure 4:**
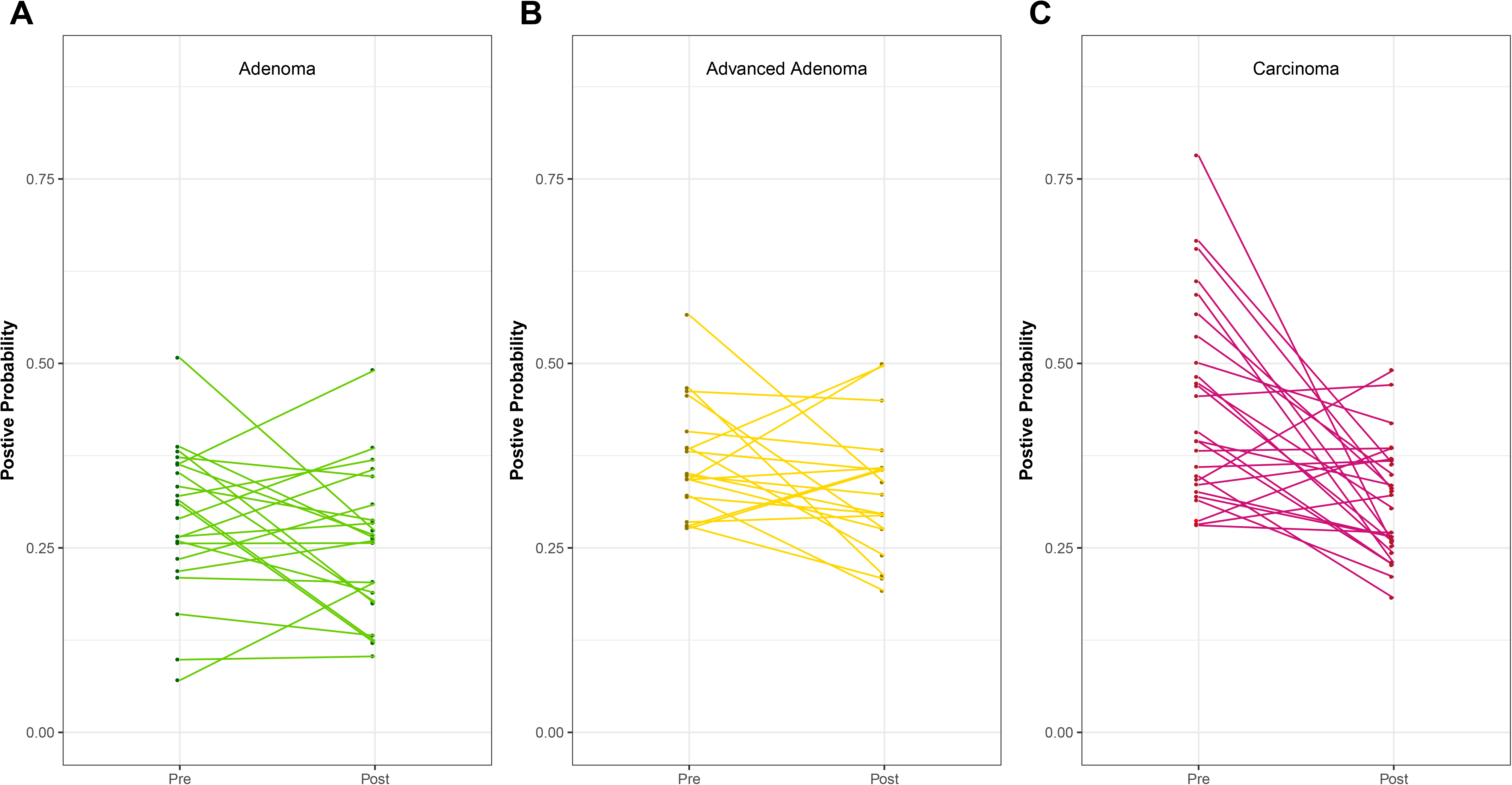
Treatment response based on models built for adenoma, advanced adenoma, or carcinoma. A) Positive probability change from initial to follow up sample in those with adenoma. B) Positive probability change from initial to follow up sample in those with advanced adenoma. C) Positive probability change from initial to follow up sample in those with carcinoma.

### Difficult to identify effects of specific treatments on the change in the microbiota

The type of treatment that the patients received varied across diagnosis groups. Those with adenomas and advanced adenomas received surgical resection (adenoma, N=4; advanced adenoma, N=4) or polyp removal during colonoscopy (adenoma, N=18; advanced adenoma, N=15) and those with carcinomas received surgical resection (N=12), surgical resection with chemotherapy (N=9), and surgical resection with chemotherapy and radiation (N=5). Regardless of treatment used there was no significant difference in the effect of these treatments on the number of observed OTUs, Shannon diversity, or Shannon evenness (P-value > 0.05). Furthermore, there was not a significant difference in the effect of the treatments on the amount of change in the community structure (P-value = 0.375). Finally, the change in the positive probability was not significantly different between any of the treatment groups (P-value = 0.375). Due to the relatively small number of samples in each treatment group, it was difficult to make a definitive statement regarding the specific type of treatment on the amount of change in the structure of the microbiota.

## Discussion

Our study focused on comparing the microbiota of patients diagnosed with adenoma, advanced adenoma, and carcinoma before and after treatment. For all three groups of patients, we observed changes in their microbiota. Some of these changes, specifically for adenoma, may be due to normal temporal variation, however, those with advanced adenoma and carcinoma clearly had large microbiota changes. After treatment, the microbiota of patients with carcinoma changed significantly more than the other groups. This change resulted in communities that more closely resembled those of patients with a normal colon. This may suggest that treatment for carcinoma is not only successful for removing the carcinoma but also at reducing the associated bacterial communities. Understanding the effect of treatment on the microbiota of those diagnosed with carcinomas may have important implications for reducing disease recurrence. It is intriguing that it may be possible to use microbiome-based biomarkers to not only predict the presence of lesions but to also assess the risk of recurrence due to these changes in the microbiota.

Patients diagnosed with adenoma and advanced adenoma, however, did not experience a shift towards a community structure that resembled those with normal colons. This may be due to the fundamental differences between the features of adenomas and advanced adenomas and carcinoma. Specifically, carcinomas may create an inflammatory milieu that would impact the structure of the community and removal of that stimulus would alter said structure. It is possible that the difference between the microbiota of patients with adenoma and advanced adenoma and those with normal colons is subtle. This is supported by the reduced ability of our models to correctly classify patients with adenomas and advanced adenomas relative to those diagnosed with carcinomas [Figure S1]. Given the irregular distribution of microbiota across patients in the different diagnosis groups, it is possible that we lacked the statistical power to adequately characterize the change in the communities following treatment.

There was a subset of patients (7 of the 26 with carcinomas) who demonstrated an elevated probability of carcinoma after treatment. This may reflect an elevated risk of recurrence. The 26.92% prevalence of increased carcinoma probability from our study is within the expected rate of recurrence (20-30% (3, 4)). We hypothesized that these individuals may have had more severe tumors; however, the tumor severity of these 7 individuals (1 with Stage I, 3 with Stage II, and 3 with Stage III) was similar to the distribution observed among the other 19 patients. We also hypothesized that we may have sampled these patients later than the rest and their communities may have reverted to a carcinoma-associated state; however, there was not a statistically significant difference in the length of time between sample collection among those whose probabilities increased (331 (246 - 358) days) or decreased (364 (301 - 434) days) (Wilcoxon Test; P-value = 0.39) (all days data displayed as median (IQR)). Finally, it is possible that these patients may not have responded to treatment as well as the other 19 patients diagnosed with carcinoma and so the microbiota may not have been impacted the same way. Again, further studies looking at the role of the microbiota in recurrence are needed to understand the dynamics following treatment.

Our final hypothesis was that the specific type of treatment altered the structure of the microbiome. The treatment to remove adenomas and advanced adenomas was either polyp removal or surgical resection whereas it was surgical resection alone or in combination with chemotherapy or with chemotherapy and radiation for individuals with carcinoma. Because chemotherapy and radiation target rapidly growing cells, these treatments would be more likely to cause a turnover of the colonic epithelium driving a more significant change in the structure of the microbiota. Although, we were able to test for an effect across these specific types of treatment, the number of patients in each treatment group was relatively small. Finally, those undergoing surgery would have received antibiotics and this may be a potential confounder. However, our pre-treatment stool samples were obtained before the surgery and the post-treatment samples were obtained long after any effects due to antibiotic administration on the microbiome would be expected to occur (344 (266 - 408) days). We also found no difference in the community structure of those that received surgery and those that did not as a treatment for adenoma or advanced adenoma.

## Conclusion

This study expands upon existing research that has established a role for the microbiota in tumorigenesis and that demonstrated the utility of microbiome-based biomarkers to predict the presence of colonic lesions. We were surprised by the lack of a consistent signal that was associated with treatment of patients with adenomas or advanced adenomas. The lack of a large effect size may be due to differences in the role of bacteria in the formation of adenomas and carcinomas or it could be due to differences in the behaviors and medications within these classes of patients. One of the most exciting of these future directions is the possibility that markers within the microbiota could be used to potentially evaluate the effect of treatment and predict recurrence for those diagnosed with carcinoma. If such an approach is effective, it might be possible to target the microbiota as part of adjuvant therapy, if the biomarkers identified play a key role in the disease process. Our data provides additional evidence on the importance of the microbiota in tumorigenesis by addressing the recovery of the microbiota after treatment and opens interesting avenues of research into how these changes may affect recurrence

## Methods

### Study Design and Patient Sampling

Sampling and design have been previously reported in Baxter, et al (24). Briefly, samples were stored on ice for at least 24h before freezing. Although we cannot exclude that this sampling protocol may have impacted the gut microbiota composition all samples were subjected to the same methodology. Study exclusion involved those who had already undergone surgery, radiation, or chemotherapy, had colorectal cancer before a baseline fecal sample could be obtained, had IBD, a known hereditary non-polyposis colorectal cancer, or familial adenomatous polyposis. Samples used to build the models for prediction were collected either prior to a colonoscopy or between one and two weeks after initial colonoscopy. The bacterial community has been shown to normalize back to a pre-colonoscopy community within this time period (27). Our study cohort consisted of 67 individuals with an initial sample as described and a follow up sample obtained between 188 - 546 days after treatment of lesion [Table 1]. Patients were diagnosed by colonoscopic examination and histopathological review of any biopsies taken. Patients were classified as having advanced adenoma if they had an adenoma greater than 1 cm, more than three adenomas of any size, or an adenoma with villous histology. This study was approved by the University of Michigan Institutional Review Board. All study participants provided informed consent and the study itself conformed to the guidelines set out by the Helsinki Declaration. The original protocol for the study did not provide for tracking patients after the follow up samples and so it was not possible for us to ascertain their diagnosis after the completion of the study.

### Treatment

For this study treatment refers specifically to the removal of a lesion with or without chemotherapy and radiation. The majority of patients undergoing treatment for adenoma or advanced adenoma were not treated surgically [Table 1] but rather via colonoscopy. All patients diagnosed with carcinomas were treated with at least surgery or a combination of surgery and chemotherapy or surgery, chemotherapy, and radiation.

The type of chemotherapy used for patients with CRC included Oxaliplatin, Levicovorin, Folfox, Xeloda, Capecitabine, Avastin, Fluorouracil, and Glucovorin. These were used individually or in combination with others depending on the patient [Table 1]. If an individual was treated with radiation they were also always treated with chemotherapy. Radiation therapy generally used 18 mV photons for treatment.

### 16S rRNA Gene Sequencing

Sequencing was completed as described by Kozich, et al. (28). DNA extraction used the 96-well Soil DNA isolation kit (MO BIO Laboratories) and an epMotion 5075 automated pipetting system (Eppendorf). The V4 variable region was amplified and the resulting product was split between four sequencing runs with normal, adenoma, and carcinoma evenly represented on each run. Each group was randomly assigned to avoid biases based on sample collection location. The pre and post-treatment samples were sequenced on the same run.

### Sequence Processing

The mothur software package (v1.37.5) was used to process the 16S rRNA gene sequences and has been previously described (28). The general workflow using mothur included merging paired-end reads into contigs, filtering for low quality contigs, aligning to the SILVA database (29), screening for chimeras using UCHIME (30), classifying with a naive Bayesian classifier using the Ribosomal Database Project (RDP)(31), and clustered into Operational Taxonomic Units (OTUs) using a 97% similarity cutoff with an average neighbor clustering algorithm (32). The number of sequences for each sample was rarefied to 10523 to minimize the impacts of uneven sampling.

### Model Building

The Random Forest (33) algorithm was used to create the three models used to classify pre and post-treatment samples by diagnosis (adenoma, advanced adenoma, or carcinoma) as well as to assess the probability that a sample was more similar to the patient’s original diagnosis or that of a disease-free patient. All models included only OTU data obtained from 16S rRNA sequencing and were processed using the caret (v6.0.76) R package. For each model we optimized the mtry hyper-parameter, which defines the number of OTUs to investigate at each split before a new division of the data was created with the Random Forest model (33). To insure that our optimization did not result in over-fitting of the data, we made 100 different 80/20 (train/test) splits of the data where the same proportion was present within both the whole data set and the 80/20 split. For each of the 100 splits, 20 repeated 10-fold cross validation was performed on the 80% component to optimize the mtry hyper-parameter by maximizing the AUC (Area Under the Curve of the Receiver Operator Characteristic). The resulting model was then tested on the 20% of the data that were held out. A summary of the mtry hyperparameter values that were tried is available in Table S5. The reported P-values for each model relative to a random labeling was assessed by comparing the distribution of the 100 80/20 splits for the correctly labeled data to the distribution of randomly labeled data.

The three diagnosis models were constructed by using the data from Baxter et al. (24), which was censored for the pre-treatment samples of the patients that we had post-treatment samples. The treatment models were then used to quantify the model probability that a patient with an initial diagnosis retained that diagnosis or a disease-free diagnosis.

### Statistical Analysis

The R software package (v3.4.1) was used for all statistical analysis. Comparisons between bacterial community structure utilized PERMANOVA (34) in the vegan package (v2.4.3). Comparisons between probabilities as well as overall differences in the median relative abundance of each OTU between pre and post-treatment samples utilized a paired Wilcoxon ranked sum test. Comparisons between different treatment for lesions utilized a Kruskal Wallis test. Where multiple comparison testing was appropriate, a Benjamini-Hochberg (BH) correction was applied (35) and a corrected P-value of less than 0.05 was considered significant. The P-values reported are those that were BH corrected. Model rank importance was determined by obtaining the median MDA from the 100, 20 repeated 10-fold cross validation and then ranking from largest to smallest MDA.

### Reproducible Methods

A detailed and reproducible description of how the data were processed and analyzed can be found at https://github.com/SchlossLab/Sze_FollowUps_Microbiome_2017. Raw sequences have been deposited into the NCBI Sequence Read Archive (SRP062005 and SRP096978) and the necessary metadata can be found at https://www.ncbi.nlm.nih.gov/Traces/study/ and searching the respective SRA study accession

## Declarations

### Ethics approval and consent to participate

The University of Michigan Institutional Review Board approved this study, and all subjects provided informed consent. This study conformed to the guidelines of the Helsinki Declaration.

### Consent for publication

Not applicable.

### Availability of data and material

A detailed and reproducible description of how the data were processed and analyzed can be found at https://github.com/SchlossLab/Sze_followUps_2017. Raw sequences have been deposited into the NCBI Sequence Read Archive (SRP062005 and SRP096978) and the necessary metadata can be found at https://www.ncbi.nlm.nih.gov/Traces/study/ and searching the respective SRA study accession.

## Authors’ contributions

All authors were involved in the conception and design of the study. MAS analyzed the data. NTB processed samples and analyzed the data. All authors interpreted the data. MAS and PDS wrote the manuscript. All authors reviewed and revised the manuscript. All authors read and approved the final manuscript.

## Acknowledgements

The authors thank the Great Lakes-New England Early Detection Research Network for providing the fecal samples that were used in this study. We would also like to thank Amanda Elmore for reviewing and correcting code error and providing feedback on manuscript drafts. We would also like to thank Nicholas Lesniak for providing feedback on manuscript drafts.

---

**Figure S1: ROC curves of the adenoma, advanced adenoma, and carcinoma models. A)** Adenoma ROC curve: The light green shaded areas represent the range of values of a 100 different 80/20 splits of the test set data and the dark green line represents the model using 100% of the data set and what was used for subsequent classification. B) Advanced Adenoma ROC curve: The light yellow shaded areas represent the range of values of a 100 different 80/20 splits of the test set data and the dark yellow line represents the model using 100% of the data set and what was used for subsequent classification. C) Carcinoma ROC curve: The light red shaded areas represent the range of values of a 100 different 80/20 splits of the test set data and the dark red line represents the model using 100% of the data set and what was used for subsequent classification.

**Figure S2: Summary of top 10% of important OTUs for the adenoma, advanced adenoma, and carcinoma models. A)** MDA of the most important variables in the adenoma model. The dark green point represents the mean and the lighter green points are the value of each of the 100 different runs. B) Summary of Important Variables in the advanced adenoma model. MDA of the most important variables in the SRN model. The dark yellow point represents the mean and the lighter yellow points are the value of each of the 100 different runs. C) MDA of the most important variables in the carcinoma model. The dark red point represents the mean and the lighter red points are the value of each of the 100 different runs.

**Figure S3: Pre and post-treatment relative abundance of CRC associated OTUs within the carcinoma model**.

## References

1. Siegel RL, Miller KD, Jemal A. 2016. Cancer statistics, 2016. CA: a cancer journal for clinicians 66:7–30.

2. Haggar FA, Boushey RP. 2009. Colorectal cancer epidemiology: Incidence, mortality, survival, and risk factors. Clinics in Colon and Rectal Surgery 22:191–197.

3. Hellinger MD, Santiago CA. 2006. Reoperation for recurrent colorectal cancer. Clinics in Colon and Rectal Surgery 19:228–236.

4. Ryuk JP, Choi G-S, Park JS, Kim HJ, Park SY, Yoon GS, Jun SH, Kwon YC. 2014. Predictive factors and the prognosis of recurrence of colorectal cancer within 2 years after curative resection. Annals of Surgical Treatment and Research 86:143–151.

5. Institute NC. SEER Cancer Stat Facts: Colon and Rectum Cancer.

6. Goodwin AC, Destefano Shields CE, Wu S, Huso DL, Wu X, Murray-Stewart TR, Hacker-Prietz A, Rabizadeh S, Woster PM, Sears CL, Casero RA. 2011. Polyamine catabolism contributes to enterotoxigenic Bacteroides fragilis-induced colon tumorigenesis. Proceedings of the National Academy of Sciences of the United States of America 108:15354–15359.

7. Abed J, Emgård JEM, Zamir G, Faroja M, Almogy G, Grenov A, Sol A, Naor R, Pikarsky E, Atlan KA, Mellul A, Chaushu S, Manson AL, Earl AM, Ou N, Brennan CA, Garrett WS, Bachrach G. 2016. Fap2 Mediates Fusobacterium nucleatum Colorectal Adenocarcinoma Enrichment by Binding to Tumor-Expressed Gal-GalNAc. Cell Host & Microbe 20:215–225.

8. Arthur JC, Gharaibeh RZ, Mühlbauer M, Perez-Chanona E, Uronis JM, McCafferty J, Fodor AA, Jobin C. 2014. Microbial genomic analysis reveals the essential role of inflammation in bacteria-induced colorectal cancer. Nature Communications 5:4724.

9. Kostic AD, Chun E, Robertson L, Glickman JN, Gallini CA, Michaud M, Clancy TE, Chung DC, Lochhead P, Hold GL, El-Omar EM, Brenner D, Fuchs CS, Meyerson M, Garrett WS. 2013. Fusobacterium nucleatum potentiates intestinal tumorigenesis and modulates the tumor-immune microenvironment. Cell Host & Microbe 14:207–215.

10. Wu S, Rhee K-J, Albesiano E, Rabizadeh S, Wu X, Yen H-R, Huso DL, Brancati FL, Wick E, McAllister F, Housseau F, Pardoll DM, Sears CL. 2009. A human colonic commensal promotes colon tumorigenesis via activation of T helper type 17 T cell responses. Nature Medicine 15:1016–1022.

11. Zackular JP, Baxter NT, Chen GY, Schloss PD. 2016. Manipulation of the Gut Microbiota Reveals Role in Colon Tumorigenesis. mSphere 1.

12. Zackular JP, Baxter NT, Iverson KD, Sadler WD, Petrosino JF, Chen GY, Schloss PD. 2013. The gut microbiome modulates colon tumorigenesis. mBio 4:e00692–00613.

13. Baxter NT, Zackular JP, Chen GY, Schloss PD. 2014. Structure of the gut microbiome following colonization with human feces determines colonic tumor burden. Microbiome 2:20.

14. Flynn KJ, Baxter NT, Schloss PD. 2016. Metabolic and Community Synergy of Oral Bacteria in Colorectal Cancer. mSphere 1.

15. Wang T, Cai G, Qiu Y, Fei N, Zhang M, Pang X, Jia W, Cai S, Zhao L. 2012. Structural segregation of gut microbiota between colorectal cancer patients and healthy volunteers. The ISME journal 6:320–329.

16. Chen H-M, Yu Y-N, Wang J-L, Lin Y-W, Kong X, Yang C-Q, Yang L, Liu Z-J, Yuan Y-Z, Liu F, Wu J-X, Zhong L, Fang D-C, Zou W, Fang J-Y. 2013. Decreased dietary fiber intake and structural alteration of gut microbiota in patients with advanced colorectal adenoma. The American Journal of Clinical Nutrition 97:1044–1052.

17. Chen W, Liu F, Ling Z, Tong X, Xiang C. 2012. Human intestinal lumen and mucosa-associated microbiota in patients with colorectal cancer. PloS One 7:e39743.

18. Shen XJ, Rawls JF, Randall T, Burcal L, Mpande CN, Jenkins N, Jovov B, Abdo Z, Sandler RS, Keku TO. 2010. Molecular characterization of mucosal adherent bacteria and associations with colorectal adenomas. Gut Microbes 1:138–147.

19. Kostic AD, Gevers D, Pedamallu CS, Michaud M, Duke F, Earl AM, Ojesina AI, Jung J, Bass AJ, Tabernero J, Baselga J, Liu C, Shivdasani RA, Ogino S, Birren BW, Huttenhower C, Garrett WS, Meyerson M. 2012. Genomic analysis identifies association of Fusobacterium with colorectal carcinoma. Genome Research 22:292–298.

20. Feng Q, Liang S, Jia H, Stadlmayr A, Tang L, Lan Z, Zhang D, Xia H, Xu X, Jie Z, Su L, Li X, Li X, Li J, Xiao L, Huber-Schönauer U, Niederseer D, Xu X, Al-Aama JY, Yang H, Wang J, Kristiansen K, Arumugam M, Tilg H, Datz C, Wang J. 2015. Gut microbiome development along the colorectal adenoma-carcinoma sequence. Nature Communications 6:6528.

21. Dejea CM, Wick EC, Hechenbleikner EM, White JR, Mark Welch JL, Rossetti BJ, Peterson SN, Snesrud EC, Borisy GG, Lazarev M, Stein E, Vadivelu J, Roslani AC, Malik AA, Wanyiri JW, Goh KL, Thevambiga I, Fu K, Wan F, Llosa N, Housseau F, Romans K, Wu X, McAllister FM, Wu S, Vogelstein B, Kinzler KW, Pardoll DM, Sears CL. 2014. Microbiota organization is a distinct feature of proximal colorectal cancers. Proceedings of the National Academy of Sciences of the United States of America 111:18321–18326.

22. Mima K, Sukawa Y, Nishihara R, Qian ZR, Yamauchi M, Inamura K, Kim SA, Masuda A, Nowak JA, Nosho K, Kostic AD, Giannakis M, Watanabe H, Bullman S, Milner DA, Harris CC, Giovannucci E, Garraway LA, Freeman GJ, Dranoff G, Chan AT, Garrett WS, Huttenhower C, Fuchs CS, Ogino S. 2015. Fusobacterium nucleatum and T Cells in Colorectal Carcinoma. JAMA oncology 1:653–661.

23. Arthur JC, Perez-Chanona E, Mühlbauer M, Tomkovich S, Uronis JM, Fan T-J, Campbell BJ, Abujamel T, Dogan B, Rogers AB, Rhodes JM, Stintzi A, Simpson KW, Hansen JJ, Keku TO, Fodor AA, Jobin C. 2012. Intestinal inflammation targets cancer-inducing activity of the microbiota. Science (New York, NY) 338:120–123.

24. Baxter NT, Ruffin MT, Rogers MAM, Schloss PD. 2016. Microbiota-based model improves the sensitivity of fecal immunochemical test for detecting colonic lesions. Genome Medicine 8:37.

25. Zeller G, Tap J, Voigt AY, Sunagawa S, Kultima JR, Costea PI, Amiot A, Böhm J, Brunetti F, Habermann N, Hercog R, Koch M, Luciani A, Mende DR, Schneider MA, Schrotz-King P, Tournigand C, Tran Van Nhieu J, Yamada T, Zimmermann J, Benes V, Kloor M, Ulrich CM, Knebel Doeberitz M von, Sobhani I, Bork P. 2014. Potential of fecal microbiota for early-stage detection of colorectal cancer. Molecular Systems Biology 10:766.

26. Zackular JP, Rogers MAM, Ruffin MT, Schloss PD. 2014. The human gut microbiome as a screening tool for colorectal cancer. Cancer Prevention Research (Philadelphia, Pa) 7:1112–1121.

27. O’Brien CL, Allison GE, Grimpen F, Pavli P. 2013. Impact of colonoscopy bowel preparation on intestinal microbiota. PloS One 8:e62815.

28. Kozich JJ, Westcott SL, Baxter NT, Highlander SK, Schloss PD. 2013. Development of a dual-index sequencing strategy and curation pipeline for analyzing amplicon sequence data on the MiSeq Illumina sequencing platform. Applied and Environmental Microbiology 79:5112–5120.

29. Pruesse E, Quast C, Knittel K, Fuchs BM, Ludwig W, Peplies J, Glöckner FO. 2007. SILVA: A comprehensive online resource for quality checked and aligned ribosomal RNA sequence data compatible with ARB. Nucleic Acids Research 35:7188–7196.

30. Edgar RC, Haas BJ, Clemente JC, Quince C, Knight R. 2011. UCHIME improves sensitivity and speed of chimera detection. Bioinformatics (Oxford, England) 27:2194–2200.

31. Wang Q, Garrity GM, Tiedje JM, Cole JR. 2007. Naive Bayesian classifier for rapid assignment of rRNA sequences into the new bacterial taxonomy. Applied and Environmental Microbiology 73:5261–5267.

32. Schloss PD, Westcott SL. 2011. Assessing and improving methods used in operational taxonomic unit-based approaches for 16S rRNA gene sequence analysis. Applied and Environmental Microbiology 77:3219–3226.

33. Breiman L. 2001. Random Forests. Machine Learning 45:5–32.

34. Anderson MJ, Walsh DCI. 2013. PERMANOVA, ANOSIM, and the Mantel test in the face of heterogeneous dispersions: What null hypothesis are you testing? Ecological Monographs 83:557–574.

35. Benjamini Y, Hochberg Y. 1995. Controlling the false discovery rate: A practical and powerful approach to multiple testing. Journal of the Royal Statistical Society Series B (Methodological) 57:289–300.

